# Skp1 Dimerization Conceals its F-box Protein Binding Site

**DOI:** 10.1101/764126

**Authors:** Hyun W. Kim, Alexander Eletsky, Karen J. Gonzalez, Hanke van der Wel, Eva-Maria Strauch, James H. Prestegard, Christopher M. West

**Affiliations:** Dept. of Biochemistry and Molecular Biology, University of Georgia, Athens, GA 30602 USA; Complex Carbohydrate Research Center, University of Georgia, Athens, GA 30602 USA; Dept. of Pharmaceutical and Biomedical Sciences, University of Georgia, Athens, GA 30602 USA; Center for Tropical and Emerging Global Diseases, University of Georgia, Athens, GA 30602 USA

**Author notes:** These authors contributed equally to this work (Hyun W. Kim, Alexander Eletsky).

## Abstract

Skp1 is an adapter that links F-box proteins to cullin-1 in the Skp1/cullin-1/F-box (SCF) protein family of E3 ubiquitin ligases that targets specific proteins for polyubiquitination and subsequent protein degradation. Skp1 from the amoebozoan *Dictyostelium* forms a stable homodimer *in vitro* with a *K*_d_ of 2.5 µM as determined by sedimentation velocity studies, yet is monomeric in crystal complexes with F-box proteins. To investigate the molecular basis for the difference, we determined the solution NMR structure of a doubly truncated Skp1 homodimer (Skp1ΔΔ). The solution structure of Skp1ΔΔ dimer reveals a 2-fold symmetry with an interface that buries ∼750 Å^2^ of predominantly hydrophobic surface. The dimer interface overlaps with subsite-1 of the F-box interaction area, explaining why only the Skp1 monomer binds F-box proteins (FBPs). To confirm the model, Rosetta was used to predict amino acid substitutions that might disrupt the dimer interface, and the F97E substitution was chosen to potentially minimize interference with F-box interactions. A nearly full-length version of Skp1 with this substitution (Skp1ΔF97E) behaved as a stable monomer at concentrations up to 500 µM and actively bound a model FBP, mammalian Fbs1, which suggests that the dimeric state is not required for Skp1 to carry out a basic biochemical function. Finally, Skp1ΔF97E is expected to serve as a monomer model for high-resolution NMR studies previously hindered by dimerization.

## INTRODUCTION

The Skp1/cullin-1/F-box (SCF) protein family of E3 ubiquitin ligases is an important mediator of protein turnover in yeast/fungi, higher plants, and animals, owing to the role of polyubiquitination in serving as a signal for recognition and degradation in the 26S-proteasome. Evidence supports the importance of the SCF complex in the protist kingdom as well,^1^ where a novel posttranslational modification has been discovered in Skp1 orthologs from groups as diverse as amoebozoa (*Dictyostelium discoideum*), apicomplexans (*Toxoplasma gondii*), and oomycetes (*Pythium ultimum*).^2^ Protist Skp1 is subject, in the presence of sufficient O_2_ and α-ketoglutarate, to hydroxylation of a Pro-residue that lies on the backside of subsite-2 of the F-box binding domain of Skp1. Once Skp1 is hydroxylated, the hydroxyproline (Hyp) residue is recognized and glycosylated by a series of glycosyltransferases, resulting in the assembly of a canonical pentasaccharide. Mutational studies show that both hydroxylation and full glycosylation are required for optimal O_2_-sensing in *Dictyostelium* and *Toxoplasma*.^1,3^ Biophysical and computational studies have generated a model by which the relatively organized structure of the pentasaccharide organizes the surrounding intrinsically disordered region of *Dictyostelium* Skp1 in such a way as to be more conducive to binding the F-box domain of FBPs,^4^ and recent studies indicate that this model is also relevant to Skp1 from *Toxoplasma*.^3^ Confirmatory biophysical studies using NMR are hampered by the dimeric state of glycosylated Skp1 (GGFGGn-Skp1) because of its relatively large size, 324 amino acids.

Recombinant guinea pig Skp1, whose sequence is identical across mammals, was previously reported to dimerize with a *K*_d_ of 1.1 µM.^5^ This value is significantly below the estimated concentration of Skp1 (a.k.a. OCP2) in the inner ear tissues (2 mM) where Skp1/OCP-2 was initially characterized, suggesting that dimerization might influence Skp1 activity in cells. Studies of *Dictyostelium* Skp1, where small angle X-ray scattering, gel filtration, and NMR studies confirmed its dimeric status at higher concentrations, indicate that glycosylation modestly inhibits dimerization.^6^ We sought to investigate the significance of Skp1 dimerization by mapping its dimer interface. The structure of Skp1 from mammals, yeast and higher plants is known when it is bound to F-box proteins.^7,8^ In contrast to the extensive sequence variations of F-box domains, the sequence and structure of Skp1 in these complexes is highly conserved. However, free Skp1 has defied structural characterization, potentially because of intrinsically disordered regions (including its C-terminal region that contributes to F-box domain recognition) that interfere with the formation of crystals for X-ray crystallography. At the same time, the Skp1 homodimer is too large for solution NMR studies without resorting to ^2^H-isotope labeling. We have found that *Dictyostelium* Skp1A remains a dimer in the absence of both an internal disordered region that was originally removed to allow crystallization with F-box proteins,^7^ and the predominantly disordered region that comprises the C-terminal subsite-2 of the F-box binding region. The doubly truncated Skp1ΔΔ dimer variant (2 × 118 aa) was sufficiently small to pursue high-resolution solution NMR structure determination using uniform ^15^N- and ^13^C-isotope labeling. We found that the homodimer interface overlaps with the F-box binding interface and confirmed the finding by Rosetta-guided mutagenesis, and discuss the implications for Skp1 function and future studies on its posttranslational regulation.

## MATERIALS AND METHODS

### Expression Plasmids

The *E. coli* expression plasmid pET19b-Skp1AΔΔ was derived from pET19b-Skp1A^9^ by site-directed mutagenesis, in which primers designed to bridge the deleted sequence as described in Figure S1C were used in a PCR reaction with Q5^®^ High-Fidelity DNA Polymerase (New England Biolabs) to amplify the modified vector. After treatment with DpnI to destroy the original vector, the sample was cloned into *E. coli* strain BL21-Gold(DE3).

For improved recovery and purification of Skp1, the *Dictyostelium* Skp1A coding sequence was codon optimized for expression in E. *coli*, and appended with an N-terminal His_6_-tag which, when excised by treatment with TEV protease, yielded the native sequence with an N-terminus of SMSL-, compared to the N-terminal SL-that occurs natively after removal of the start Met.^10^ The cDNA (Figure S1A) was synthesized and provided in pUC57 by GenScript, excised using NcoI and BamHI, and ligated into the NcoI and BamHI restriction sites of pET19b, yielding pET19b-His_6_DdSkp1A-optim. A second cDNA in which 12 internal amino acids (SPQGDDKKDEKR) were replaced with GGSG (Figure S1B) was synthesized and similarly ligated into pET19B yielding pET19b-His_6_DdSkp1AΔLoop-optim. This plasmid was modified to generate an F97E point mutation by site-directed mutagenesis, in which the indicated primers were used in a PCR reaction to amplify the modified vector as above.

### Expression and Purification of Skp1 constructs

Skp1 and Skp1ΔΔ, which each lacked an affinity tag to ensure native-like behavior, were purified from *E. coli* to near homogeneity under non-denaturing conditions (DEAE, phenyl, Q and S200 Superdex columns) as described previously.^6^ Sample purity and integrity was assessed by SDS-PAGE and Coomassie blue staining to be >90%.

*E. coli* cells expressing His_6_Skp1Δ or His_6_Skp1ΔF97E were incubated at 37 °C in 2 × 1 L of Terrific Broth medium in the presence of 100 µg/ml ampicillin. At an OD_600_ of 0.6, protein expression was induced by addition of 125 µM isopropyl 1-thio-β-D-galactopyranoside (IPTG) at 20 °C. After 12-16 h, bacteria were collected by centrifugation at 5000 × *g* for 10 min and resuspended in 50 mM Na^+^/K^+^ phosphate (pH 7.8), 300 mM NaCl, 5 µg/ml aprotinin, 5 µg/ml leupeptin at 4 °C. Cells were lysed using a probe sonicator (model 500, Thermo Fisher Scientific) for a total sonication time of 5 min. The lysate was centrifuged at 25,000 × *g* for 45 min at 4 °C, and the supernatant was immediately applied to a 1.5-ml column of Co^2+^ Talon resin (Clontech) pre-equilibrated at 4 °C in the buffer described above. The column was washed successively with the same buffer supplemented with either 1 M NaCl, 10% glycerol, or 5 mM imidazole. Protein was eluted with buffer containing 300 mM imidazole, and dialyzed against 50 mM Tris-HCl (pH 8.0), 300 mM NaCl, 1 mM EDTA, 2 mM β-mercaptoethanol. The sample was incubated overnight at 22 °C with His_6_TEV protease to cleave the His_6_-tag from Skp1, and the sample was re-applied to the Talon resin. The flow-through was concentrated to 1.5 ml using a spin concentrator (Amicon) with a 3 kDa molecular weight cut-off. The concentrated sample was further purified over a Superdex 200 Hi-load 16/60 gel filtration column (GE Healthcare) pre-equilibrated with 20 mM potassium phosphate (pH 7.4), 50 mM KCl, and 0.5 mM TCEP. The sample was estimated to be >95% pure by SDS-PAGE and staining with Coomassie blue.

Stable isotope labeled Skp1ΔΔ and His_6_Skp1ΔF97E were prepared by expression in *E. coli* in the presence of isotope enriched minimal media as previously described.^9^ His_6_Skp1ΔF97E was uniformly enriched with ^15^N, and Skp1ΔΔ was labeled with ^15^N and ^13^C. The final Skp1ΔΔ NMR sample (105 µl in a 3-mm Shigemi tube) contained a 1:1 mixture of ^15^N,^13^C-Skp1ΔΔ and natural abundance Skp1ΔΔ at ∼1.0 mM concentration in 20 mM MES-NaOH (pH 6.0), 50 mM NaCl, 5 mM dithiothreitol, 0.05% NaN_3_. ^15^N-labeled Skp1ΔF97E NMR samples (300 µl in 5-mm Shigemi tubes) were prepared at concentrations of 100 µM and 500 µM in the same buffer. All NMR samples contained 10% D_2_O for spectrometer lock.

### Analytical Ultracentrifugation

Protein was quantified based on molar absorptivity calculated from the protein sequence using ProtParam.^11^ Samples were loaded into 12-mm double-sector Epon centerpieces equipped with quartz windows and equilibrated for 2 h at 20 °C in an An60 Ti rotor. Sedimentation velocity data were collected using an Optima XLA analytical ultracentrifuge (Beckman Coulter) at 50,000 rpm at 20 °C. Data were recorded with absorbance optics at 280 nm, 230 nm or 215 nm in radial step sizes of 0.003 cm. SEDNTERP^12^ was used to model the partial specific volume as well as the density and viscosity of the buffer. SEDFIT^13^ was used to analyze sedimentation data. All data were modeled as continuous *c(s)* distributions and were fit using baseline, meniscus, frictional coefficient, and systematic time-invariant and radial-invariant noise. Predicted sedimentation coefficient (s) values for Skp1 monomer and dimer states were calculated using HYDROPRO^14^ with a homology model generated on the ROBETTA server.^15^ Data fit and *c(s)* plots were generated using GUSSI.^16^ Weight-averaged S values (S_w_) at each concentration were determined by integrating *c(s)* distributions. Constructed S_w_ isotherms were fitted with a A+A⇌AA self-association model using SEDPHAT.^17^

### NMR Spectroscopy and Structure Determination of Skp1ΔΔ

NMR spectra for Skp1ΔΔ were acquired at 35 °C using a Bruker AVANCE NEO 800 MHz spectrometer equipped with a 5-mm cryogenic TCI ^1^H{^13^C,^15^N} probe, and an Agilent VNMRS 600 MHz spectrometer equipped with a 3-mm cryogenic ^1^H{^13^C,^15^N} probe. NMR spectra for 100 µM and 500 µM Skp1ΔF97E samples were acquired using a Bruker AVANCE NEO 900 MHz spectrometer equipped with a 5-mm cryogenic TXO ^13^C,^15^N{^1^H} probe, and the 600 MHz spectrometer equipped with a 5-mm cryogenic ^1^H{^13^C,^15^N} probe. The acquired NMR spectra are summarized in Table S1. NOE mixing times were 70 ms for ^13^C/^15^N-edited [^1^H,^1^H]-NOESY and 120 ms for ^13^C/^15^N-filtered ^13^C/^15^N-edited [^1^H,^1^H]-NOESY experiments. Fourier transform was performed with TopSpin (Bruker BioSpin) for Bruker NMR data, and NMRPipe^18^ for Varian NMR data. ^1^H chemical shifts were referenced relative to 4,4-dimethyl-4-silapentane-1-sulfonic acid (DSS), and ^13^C and ^15^N chemical shifts were referenced indirectly via gyromagnetic ratios. 2D and 3D NMR spectra were analyzed using CARA.^19^

Relaxation delays were 0.1, 0.2, 0.3, 0.4, 0.7, 1.0, 1.5, and 2.0 seconds in 1D proton-detected ^15^N T_1_ experiments, and 10, 30, 50, 70, 90, 110, 130, 150 and 170 ms in 1D proton-detected ^15^N T_2_ experiments acquired for the 500 µM Skp1ΔF97E sample. 1D ^15^N T_1_/T_2_ relaxation spectra were processed and analyzed with VnmrJ v4.2 (Agilent Inc). Macro “tc” (wiki.nesg.org) was used to integrate the regions between ^1^H chemical shifts of 8.8 and 9.9 ppm, determine average ^15^N T_1_ and T_2_ relaxation times via exponential fitting, and calculate global rotational correlation time τ_c_.

Sequence-specific backbone and side-chain resonance assignments for Skp1ΔΔ were derived using CARA based on existing resonance assignments of full-length Skp1.^9^ Structure calculation of the Skp1ΔΔ homodimer was performed using CYANA^20^ based on ^1^H-^1^H upper distance constraints derived from 13C/^15^N-edited [^1^H,^1^H]-NOESY, as well as backbone φ and ψ and side-chain χ_1_ dihedral angle restraints from TALOS-N.^21^ Automated NOESY peak assignment was performed initially with CYANA, with 22 manually assigned intermolecular 5 Å ^1^H-^1^H upper distance constraints (Table S2) applied after cycle 1 of simulated annealing. These intermolecular ^1^H-^1^H upper distance constraints were derived from selected strong peaks in a ^13^C/^15^N-filtered ^13^C/^15^N-edited [^1^H,^1^H]-NOESY spectrum. Only those peaks that could be unambiguously assigned and could not be explained by intramolecular contacts were chosen. After several rounds of iterative refinement of NOE peak assignments and calibration of distance constraints, the final structure calculation was performed with CYANA. Stereospecific resonance assignment of methylene proton spins and methyl groups of Leu and Val residues were obtained with the GLOMSA module of CYANA. Out of 100 calculated conformers, 20 conformers with the lowest target function values were selected for subsequent refinement in explicit water bath using the program CNS^22^ with upper distance constraints relaxed by 5%. The structure statistics are outlined in Table 1.

**Table 1.** Skp1ΔΔ dimer NMR structure statistics (PDB ID: 6V88, BMRB ID:30696)

|  |  |
| --- | --- |
| **Completeness of resonance assignments<sup>a</sup> [%]** |  |
| *Backbone/Side-chain* | 100.0/100.0 |
| **Conformation-restricting distance constraints<sup>b</sup>** |  |
| *Intra-residue* [$i = j$] | 696 |
| *Sequential* [$\vert i - j\vert = 1$] | 1256 |
| *Medium range* [$1 < \vert i - j\vert < 5$] | 1360 |
| *Long range* [$\vert i - j\vert \geq 5$] | 1470 |
| *Total* | 4782 |
| *Intermolecular NOE constraints (included in above)* | 182 |
| *Dihedral angle constraints* | 406 |
| **NOE constraints per restrained residue (of those, long range)** | 22.6 (6.4) |
| **CYANA target function [$\text{\AA}^2$]** | 2.33 |
| **Average number of distance constraint violations per conformer** |  |
| *0.1 - 0.2 $\text{\AA}$* | 9.25 |
| *0.2 - 0.5 $\text{\AA}$* | 2.2 |
| *>0.5 $\text{\AA}$* | 0 |
| **Average number of dihedral angle constraint violations per conformer** |  |
| *>10°* | 0.1 |
| **Average RMSD from mean coordinates [$\text{\AA}$]** |  |
| *backbone atoms<sup>c</sup> (all)* | 0.8 (1.2) |
| *heavy atoms<sup>c</sup> (all)* | 1.1 (1.4) |
| **Global quality scores<sup>c</sup> (raw / Z-score)** |  |
| *PROCHECK<sup>2</sup> G-factor (phi-psi)* | 0.16/0.94 |
| *PROCHECK<sup>2</sup> G-factor (all)* | 0.14/0.83 |
| *Molprobity<sup>3</sup> clash score* | 2.55/1.09 |
| *ProsaII<sup>4</sup>* | 0.74/0.37 |
| **Molprobity<sup>3</sup> Ramachandran summary [%]** |  |
| *Most favored regions* | 98.7 |
| *Additionally allowed regions* | 1.3 |
| *Disallowed regions* | 0.0 |

<sup>a</sup> Commonly observed protein NMR resonances. Excludes amino group of N-terminal Ser, side-chain amino groups of Lys, side-chain guanidinium groups of Arg, carboxyl groups of Asp and Glu, thiol and hydroxyl $^1\text{H}$ of Cys, Ser, Thr and Tyr, and non-protonated aromatic $^{13}\text{C}$ .
<sup>b</sup> Calculated with Protein Structure Validation Software (PSVS 1.5; http://psvs.nesg.org/)
<sup>c</sup> Ordered residue ranges: 3-34, 38-44, 46-64, 74-115
<sup>1</sup> Ref. 32
<sup>2</sup> Ref. 33
<sup>3</sup> Ref. 34
<sup>4</sup> Ref. 35

Predicted rotational correlation time τ_c_ was calculated for Skp1ΔΔ and Skp1ΔF97E using HYDRONMR,^23^ assuming water viscosity as 0.0072 cP at 35 °C. The lowest energy conformer of Skp1ΔΔ NMR ensemble was used, and only the first chain was used to calculate τ_c_ of a hypothetical Skp1ΔΔ. Homology model of Skp1ΔF97E monomer for τ_c_ calculation was generated with SWISS-MODEL based on X-ray structure of human Skp1 (PDB ID 3L2O) as a template. A hypothetical dimer model of Skp1ΔF97E was then produced by structural alignment of individual subunits to those of Skp1ΔΔ dimer in Chimera (http://www.rbvi.ucsf.edu/chimera),^24^ followed by adjustment of backbone dihedral angles to eliminate clashes in the C-terminal region and energy minimization.

### Analysis of the Dimer Interface

Computational investigation of the dimer interface was performed using the conformer with the lowest CYANA target function of the initial NMR structures of Skp1ΔΔ. To prevent inaccurate predictions due to small clashes in the structure, the protein was prepared using the standard Rosetta optimization protocol, “FastRelax”.^25,26^ Briefly, five cycles of rotamer packing and minimization were carried out, ramping up the repulsive weight in the scoring function within each cycle. After three rounds of symmetrical FastRelax with atom-atom pair distance constraints, the quality of the generated models was validated with the Molprobity web service.^27^ The lowest scoring structure based on the Rosetta energy score and the Molprobity score was selected for mutational analysis.

To identify mutations that could disrupt the Skp1 dimerization, an all amino acids-scanning mutagenesis was carried out across the homodimer interface *in silico*. The “flex ddG” protocol implemented in RosettaScripts^28,29^ was used to model and predict the effect of the mutations on the binding free energy of the complex. All parameters in the protocol were set up according to the default values described previously.^28^ Overall, the flex ddG method takes advantage of the Rosetta backrub approach^30^ to sample side-chain and backbone conformational changes around the mutated position. Once the backrub ensembles are generated, the structures are optimized by sidechain repacking and torsion minimization. The interface ΔΔG score corresponds to the average difference in binding free energy between the mutant structure and the wild-type complex. Stabilizing mutations are defined as those with interface ΔΔG scores < −1.0 Rosetta energy units (REU), while destabilizing mutations are assigned to interface ΔΔG scores > 1.0.^28^

To determine which dimer-destabilizing substitutions do not perturb the stability of the individual monomers, the change in the total free energy of the monomer was estimated for all mutations. This analysis was performed with the current state-of-the-art Rosetta ΔΔG protocol, “cartesian_ddg”.^31^ Prior to the simulations, the refined wild-type monomer was relaxed in Cartesian space, constraining backbone and sidechain coordinates. The model with the lowest Rosetta score was then used as input for the cartesian_ddg protocol. Within the cartesian_ddg application, the protein was relaxed again in the Cartesian space, allowing movement of only the backbone and sidechains around the mutated position.^31^ All parameters in the method were configured as previously described.^31^ The total ΔΔG score was finally considered as the difference in the total ΔΔG between the mutant and the wild-type monomer, multiplied by an energy scaling factor of 1.0/2.94. As above, stabilizing mutations correspond to total ΔΔG scores < −1.0, and destabilizing mutations refer to total ΔΔG scores > 1.0.

All Rosetta commands for this report were run with the same Rosetta static executable (RosettaCommons/main.git2019-03-07, version 4ab48a76160c888257155619edb9817845bd8a67). The protocols previously described can be found at Supplementary information-Scripts.

### Analytical Gel Filtration

Skp1Δ and Skp1ΔF97E with or without Fbs1 at a limiting concentration were subjected to Superdex 200 PC 3.2/30 gel filtration analysis using a Pharmacia SMARTSystem HPLC as previously described.^6^

## RESULTS

### Characterization of the Skp1 Dimer

Sedimentation velocity experiments were conducted on *Dictyostelium* Skp1A (Skp1) that was recombinantly expressed without a peptide tag in *E. coli*. Over a concentration range of 0.5 – 45 µM, the samples yielded peaks at 1.8 S and 2.7 S (Figure 1A), values which are slightly less than the predicted S-values for monomer and dimer forms, 1.9 S and 2.8 S. The homology model used for predicting S-values assumed that the C-terminal region of Skp1 is organized as α-helices as occurs in complexes with F-box proteins. However, the C-terminal region of free Skp1 is predominantly disordered based on previous NMR studies,^9^ which is expected to cause Skp1 to sediment more slowly and would explain the slight discrepancy between the observed and predicted S-values. The separate peaks indicate that interconversion between monomer and dimer states is slow relative to the time scale of sedimentation. A Skp1 dimer binding isotherm constructed using weighted S-values from across the concentration range yielded a dissociation constant for the Skp1 dimer of 2.5 µM (Figure 1B).

**Figure 1.**
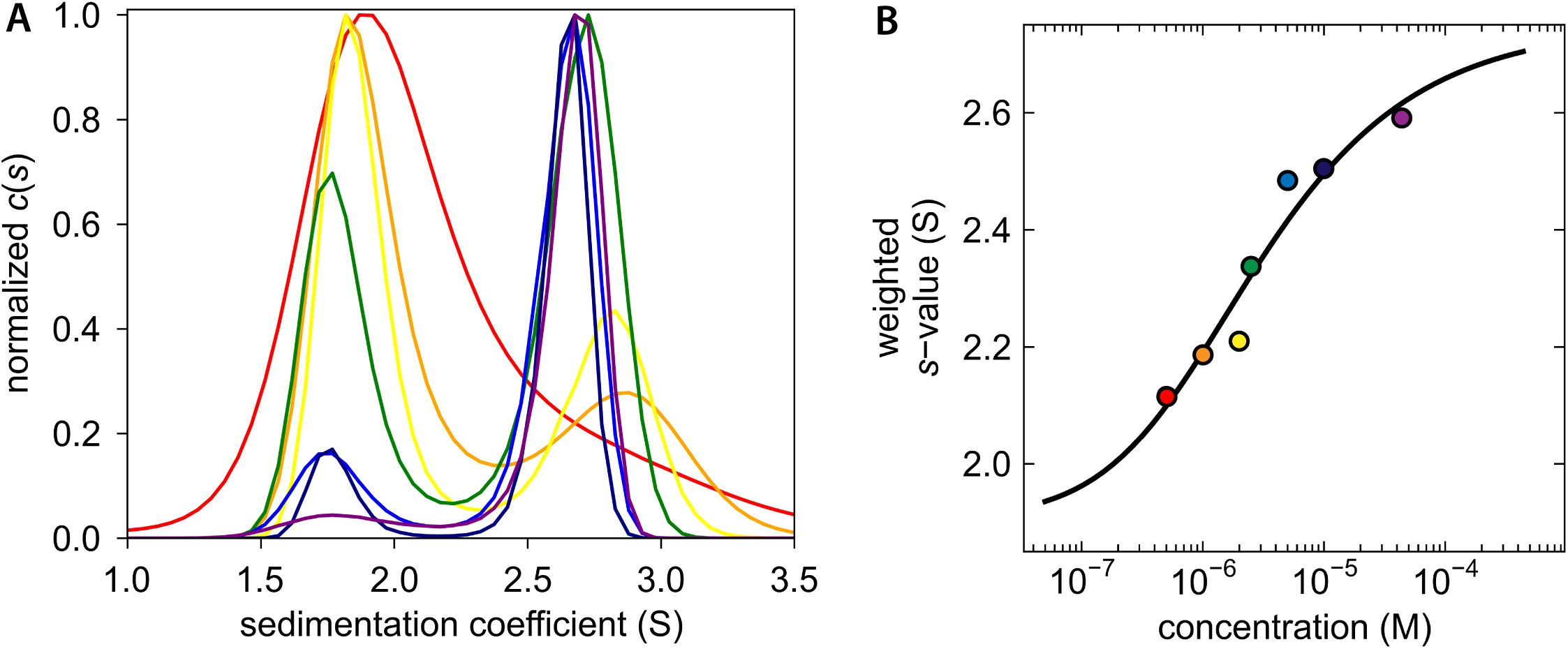
Sedimentation velocity analysis of *Dd*Skp1. (A) *c(s)* distribution reveals concentration dependence of dimerization. The concentration range is depicted by a rainbow spectrum with the lowest concentration in red and the highest in purple. (B) An isotherm was constructed with weighted s-values (S_w_); the fitted model indicates a *K*_d_ of 2.5 µM. The color of each data point corresponds to the respective *c(s)* distribution in panel A.

Skp1 has resisted crystallization and the large size of the full-length homodimer (324 amino acids) inhibited structure determination by NMR. To initiate mapping of the dimer interface, we examined a truncated Skp1 variant, Skp1ΔΔ, which lacks the mainly disordered C-terminal F-box binding domain and an internal disordered loop that is frequently removed for Skp1/FBP crystallization (Figure 2A). We demonstrated that Skp1ΔΔ (118 × 2 amino acids) still forms a stable homodimer based on sedimentation velocity studies, which yielded an S-value of 1.9, in agreement with the 1.9 S-value predicted by HYDROPRO (Figure S2). Also, a 2D [^15^N,^1^H] HSQC spectrum of Skp1ΔΔ correlated well with 2D [^15^N,^1^H] TROSY of full-length Skp1 (data not shown), indicating that truncations in Skp1ΔΔ do not perturb the overall structure.

**Figure 2.**
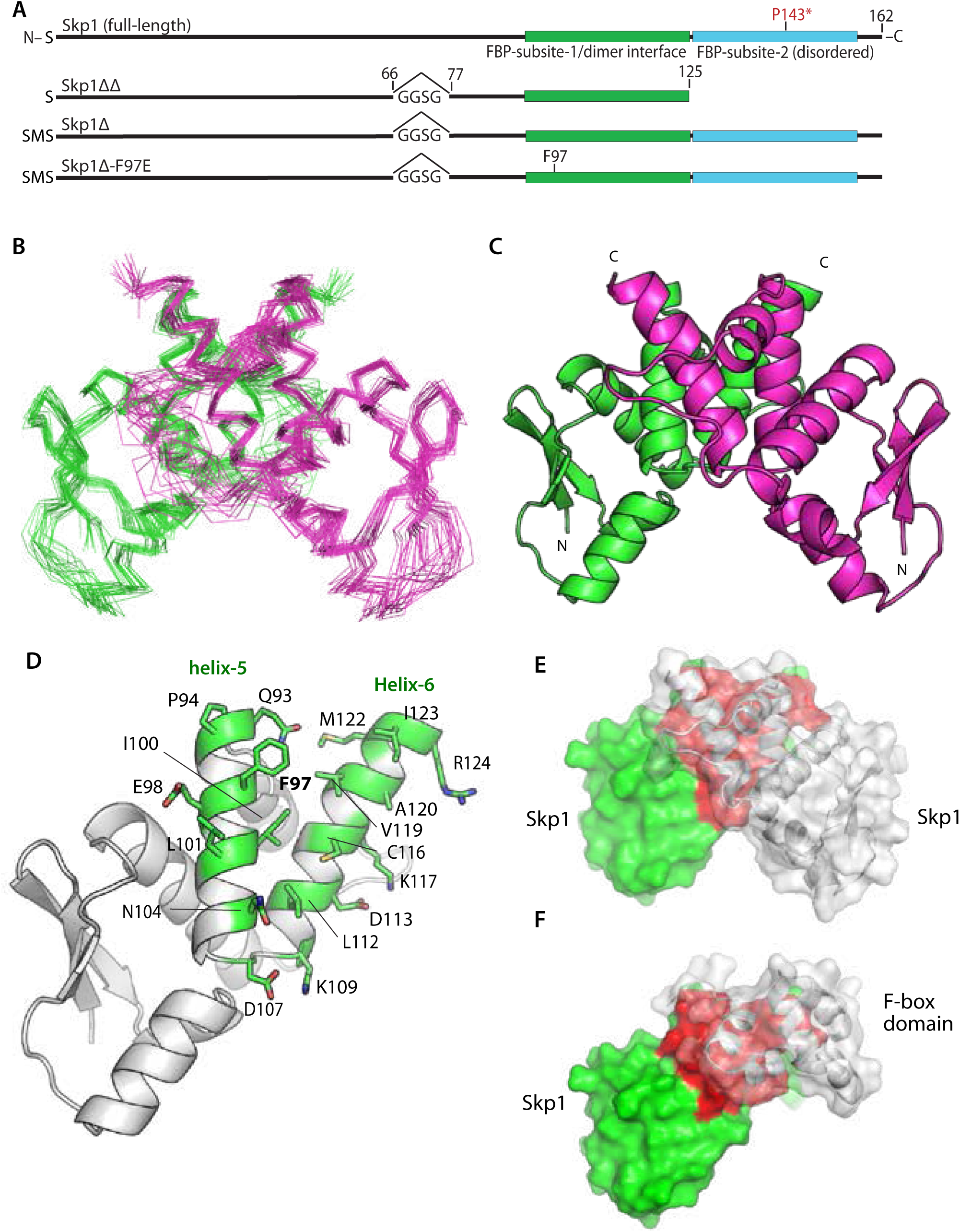
Structure of the Skp1 dimer. (A) Domain diagrams of the constructs examined. Note that versions derived from His_6_Skp1 have a SerMet-extension beyond the native Ser-resulting from Met removal. See Figure S1 for details. (B) Superimposition of Cα-traces of 20 calculated conformers of Skp1ΔΔ. (C) Ribbon representation of the lowest energy Skp1ΔΔ conformer (PDB ID 6V88). Dimer subunits are colored in green or magenta. A 2-fold axis of rotational symmetry lies vertically between the subunits. (D) Ribbon representation of a single Skp1ΔΔ, with the residues contributing to intermolecular contacts (<5 Å) shown in green with stick representations of their side chains. (E) Surface representation of the Skp1ΔΔ dimer is shown with the rear subunit colored in green and red, and the front in transparent gray. Red shading represents the homodimer contact region. (F) Surface representation of a hypothetical Skp1ΔΔ/F-box heterodimer model, generated by substitution of a single Skp1ΔΔ subunit for Skp1 in a human Skp1/FBXW7 complex (PDB ID 5V4B). Coloration is as in E, with FBXW7 residues 2263-2355 in gray.

### Solution NMR Structure of Skp1ΔΔ

Using a suite of standard NMR experiments (Table S1), we obtained complete sequence-specific assignments of backbone and side-chain ^1^H, ^15^N and ^13^C resonances of Skp1ΔΔ (Table 1; Figure S3). These resonance assignments allowed us to pursue structure calculations based on ^1^H-^1^H distance constraints derived from 3D ^15^N- and ^13^C-edited [^1^H,^1^H] NOESY spectrum. To ensure proper modeling of the subunit interaction, we applied weak intermolecular distance ^1^H-^1^H constraints (Table S2) derived from NOE peaks identified in a separate ^13^C/^15^N-filtered and ^13^C-edited [^1^H,^1^H] NOESY spectrum recorded with a sample of mixed U-^15^N,^13^C-labeled and natural abundance Skp1ΔΔ. Representative fragments of NOESY spectra are shown in Figure S4. Full attenuation of intra-chain NOE cross-peaks was not achieved in the ^13^C/^15^N-filtered spectrum (right strip in Figure S4) because of incomplete isotope incorporation (∼85%). However, comparison with the corresponding strip from the NOESY spectrum without isotope filtering (strip on the left) allowed us to distinguish between peaks of comparable intensity (inter-chain) and peaks with significantly reduced intensity in the filtered set (intra-chain).

We obtained a high-quality solution NMR structure of Skp1ΔΔ (Table 1; Figure 2). A ribbon diagram of the best scoring structure of the top 20 conformers (Figure 2B) is shown in Figure 2C. The structure features a semi-parallel orientation of subunits with respect to their N- and C-termini, in contrast to the previously hypothesized dimerization model from SAXS analysis.^6^ The dimer interface is organized as a four-helix bundle, with symmetrical packing contributions from residues of helices 5 & 6 of each chain (Figure S5) as identified using PISA.^36^ The interface buries ∼743 Å^2^ of predominantly hydrophobic surface whose participating amino acids are labeled in Figure 2D. Superimposition of the corresponding Cα atoms with human Skp1 from a crystal structure in complex with the human FBP βTRCP revealed an RMSD of 1.1 Å (Figure S6), indicating that the structure of amino acids 1-125 of free Skp1 changes little when complexed as a monomer with FBPs. Furthermore, the homodimer interface involves the previously described subsite-1 of the binding site for F-box domains.^7^ The physical overlap (compare Figures 2E, F) was quantified by modeling the F-box domain of human FBXW7 from a crystal structure with human Skp1, with Skp1ΔΔ. This revealed a ∼650 Å^2^ overlap of the homodimer and heterodimer interfaces, in the region of helices 5 and 6, and explains why only the Skp1 monomer is found in complexes with FBPs.

### Conservation of the Dimer Interface

The two α-helices contributing to the homodimer interface, extending from L96 to I123, are highly conserved throughout phylogeny. Furthermore, each dimer contact residue except K117 is almost perfectly conserved from stramenopiles to humans (Figure S8), and the region is immediately surrounded by Gly or Pro residues and length variations (indels), indicating that this region represents a functional unit under selective pressure to remain intact. This is consistent with the finding that human Skp1 also dimerizes in this concentration range.^5^

### Computationally-Guided Selection of a Skp1 Monomer Mutant

To test the dimer structure model, we searched for point mutations of interface amino acids that might destabilize the dimer. An initial alanine-scanning mutagenesis calculation suggested Phe97 Leu101, and Ile123 as potential destabilizing positions based on a weakening of the binding free energy of the complex (interface ΔΔG score > 2) (Figures 3A, 3B). All 20 amino acids were then substituted at each position to predict the effect of different side chains. Indeed, most substitutions at these 3 positions were destabilizing, and 85% of the variations at position 97 yielded interface ΔΔG scores greater than 1.5. The mutations F97G, F97D and F97E showed the highest dimer-destabilizing effect, with interface ΔΔG scores of 5.41, 5.23 and 4.87, respectively (Figure 3C).

**Figure 3.**
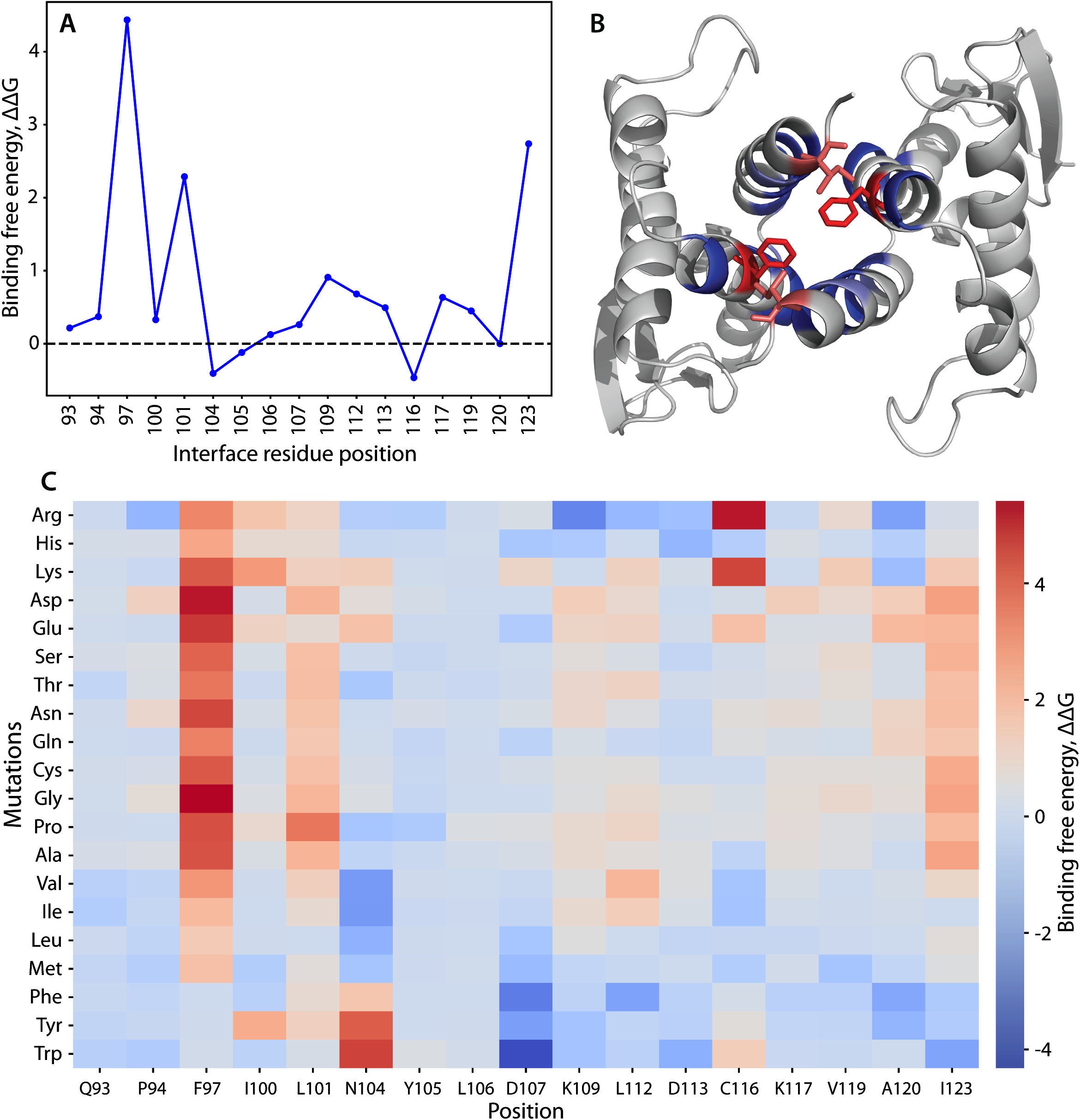
Computational scanning mutagenesis of the Skp1ΔΔ homodimer interface. (A) Alanine-scanning mutagenesis using Rosetta. Changes in binding free energy upon replacement with alanine are shown according to the interface residue positions. (B) Skp1 dimer protein-protein interface. Residues with the highest binding free energy change (Phe97 and Ile123) are emphasized in stick representation and in red; other mutated residues are in blue. See panel C for color code explanation. (C) Heatmap of the changes in binding free energy upon all amino acid substitutions. Effects of amino acid replacements are shown for each interface position. The colors represent the changes in the binding free energy of the dimer (interface ΔΔG score). Values greater than one (warmer colors) indicate destabilizing mutations, and values less than one (colder colors) imply stabilizing mutation.^28^ Compare with effects on the monomer state (Fig. S7).

To determine whether the dimer-destabilizing mutations might affect monomer stability, the total monomer ΔΔG was estimated for each substitution. Remarkably, most of the mutations at positions 97, 101, and 123 presented total ΔΔG scores between 1 and −1 (Figure S7). In particular, the highly dimer destabilizing F97E substitution is predicted to have a neutral effect on monomer stability with its ΔΔG score at 0.15.

Based on the above analysis, the conserved Phe97 was changed to Glu in the nearly full length Skp1Δ isoform. Glu was chosen over Asp to allow extension of the carboxyl group to provide better accessibility to solvent rather than clashing when binding an FBP. As predicted by the dimer interface model, Skp1ΔF97E eluted as a monomer based on gel filtration and was predominantly a monomer by AUC at 100 µM (Figure 4A). A 2D [^1^H,^15^N]-HSQC spectrum recorded for a ^15^N-labeled Skp1ΔF97E sample exhibited peak dispersion consistent with a well-folded protein (Figure 4B). The peak pattern was comparable to that of wild-type Skp1 considering that there are a number of differences including an N-terminal SM-extension, replacement of 12 amino acids with 4 different amino acids in the internal loop, and truncation after amino acid 125 (not shown). Based on average ^15^N T_1_ and T_2_ relaxation times measured for all ^1^H amide resonances between 8.8 and 9.9 ppm in a 500 µM sample of Skp1ΔF97E, the effective rotational correlation time (*τ*_c_) was calculated at 9.4 ± 0.5 ns. This is close to the 10.9 ns value predicted for the monomer (Table 2), and contrasts with the previously reported 19.9 ± 2.2 ns value of the full-length protein.^4^ Thus, Skp1ΔF97E remains predominantly monomeric even at high concentrations typical for solution NMR.

**Table 2.** Predicted and Experimental τ_c_ for Skp1 isoforms at 35°C.

| | Predicted<sup>a</sup> $\tau_c$ (ns), monomer/dimer | Experimental $\tau_c$ (ns) |
| --- | --- | --- |
| Skp1ΔF97E, 500 μM | 10.9/19.6 | $9.4 \pm 0.5$ |
| Skp1(native), 850 μM | Similar to above | $19.5 \pm 2.2^b$ |

<sup>a</sup> Predicted using HYDRONMR<sup>23</sup>
<sup>b</sup> from Ref. 7

**Figure 4.**
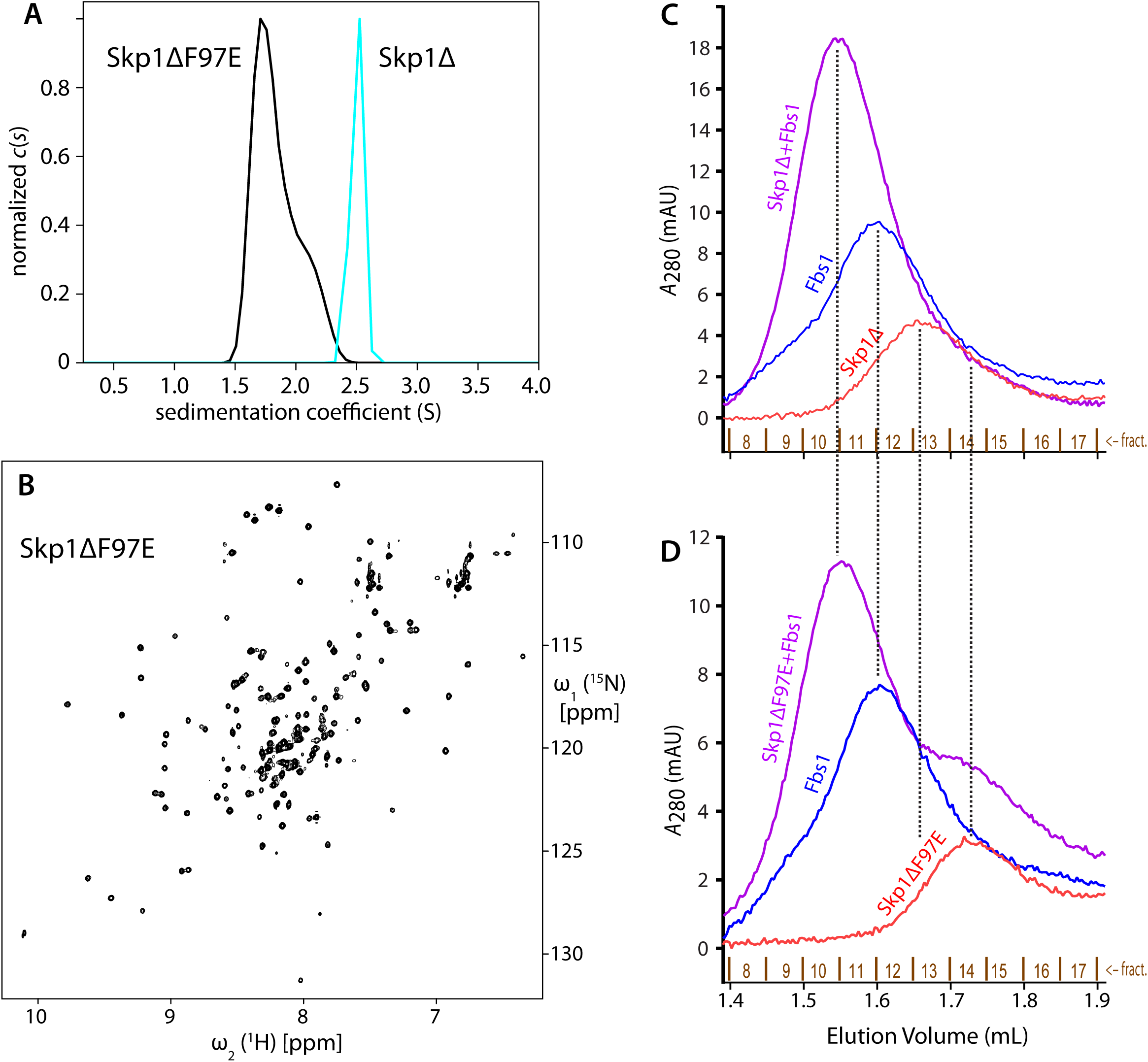
Skp1ΔF97E is a stable and functional monomer in solution. (A) *c(s)* distributions of 100 µM Skp1Δ or Skp1ΔF97E are shown in cyan and black, respectively. (B) ^1^H/^15^N-HSQC of 100 µM Skp1ΔF97E at 900 MHz and 35°C, with a 4 h collection time. The 500 μM spectrum (not shown) was indistinguishable. (C, D) Skp1ΔF97E binds the model F-box protein Fbs1. His_6_Fbs1 (1.5 μM) and an estimated 2.25 µM Skp1Δ (C) or Skp1Δ(F97E) (D) were analyzed on a Superdex 200 gel filtration column. Elution was monitored by *A*_280_, which favors detection of Fbs1 relative to Skp1 because of its higher extinction coefficient.

### Skp1ΔF97E is Binding Competent with a Model FBP

Phe97 is conserved as a Phe or Tyr in known Skp1s (Figure S8). Analysis of Skp1 in crystal structures of complexes with 3 different FBPs (Tir1, βTRCP1, Fbs1) shows that, compared to the homodimer, Phe (or Tyr) resides in a different rotamer state with solvent exposure (not shown). This suggests that the F97E replacement will, though removing a favorable hydrophobic contact, not directly disrupt the FBP interaction. To test the functionality of Skp1ΔF97E, we used a Superdex200 gel filtration column to examine the elution profile of Skp1 in the absence and presence of a heterologous FBP, guinea pig Fbs1. As previously described^6^ and replicated in Figure 4C, a mixture of Fbs1 and Skp1 eluted prior to the elution positions of either protein alone. As shown in Figure 4D, Skp1ΔF97E exhibited similar behavior. In addition, Skp1ΔF97E clearly eluted later than native Skp1, consistent with its monomeric character as described by AUC and its NMR-derived rotational correlation time. Thus Skp1(F97E) retained its FBP binding function, though a possible reduction in affinity is not excluded by this analysis.

## DISCUSSION

Our findings confirm that dimerization is a highly conserved property of Skp1, based on similar dissociation constant values from organisms as phylogenetically distant as *Dictyostelium* and humans. Their measured *K*_d_ values range from 1.1 to 2.5 µM under *in vitro* conditions, though actual affinities in the cell may vary. These values are similar to the predicted total monomer Skp1 concentration in a mammalian cell line, ∼2 µM.^37^ The significance of dimerization is suggested by the exceptionally high degree of conservation of the contact residues (labeled D in Figure S8), and its ability to mask the hydrophobic character of this solvent exposed surface.

The Skp1 dimer interface occupies ∼740 Å^2^ of predominantly hydrophobic surface, a substantial area that overlaps subsite-1 of its FBP binding site (Figure 2E). The corresponding region in human Skp1 (residues 99-130 in human vs. residues 93-124 in *Dictyostelium* Skp1, Figure S8) exhibited a paucity of NMR resonance assignments in a recent study of human Skp1,^38^ indicating potential effects of dimerization in human Skp1. The homodimerization model was supported by the predicted effect of an amino acid substitution within the interface, F97E, to destabilize the interaction (Figure 4A) without unfolding the protein (Figure 4B). Indeed, monomeric Skp1 maintained its ability to bind a model FBP, Fbs1 (Figure 4C, D), and was also competent to be enzymatically hydroxylated and fully glycosylated *in vitro* (unpublished data). A further contribution to the dimer interface from beyond the truncation site at residue 125 seems unlikely, because previous studies indicated substantial disorder for this region in the free dimer.^9^

Based on the new structure, the dimer interface also contributes to subsite-1 of the F-box binding region of Skp1, which explains why Skp1 is a monomer in complexes with FBPs.^8^ Interestingly, 8 of the 13 alleles of Skp1 known to affect its function in budding and fission yeast have point mutations located on this region,^39-43^ raising the question of whether these mutations affect dimerization, FBP binding, or both. Since the Skp1/FBP interaction is very strong (*K*_d_ for binding to the FBP Fbs1/OCP1 is ca. two orders of magnitude smaller than the homodimerization *K*_d_)^44^, and quantitative mass spectrometry indicates that Skp1 is not in great excess over FBPs in cells,^37^ the average concentration of Skp1 does not appear to be high enough to generate a substantial homodimer pool. However, if higher local concentrations occur in the cell, homodimerization might occur to protect Skp1 from interacting promiscuously with other macromolecules. Our ability to selectively perturb dimerization relative to Fbs1 binding using the F97E mutation might allow an investigation of this question *in vivo*; however, we cannot exclude the possibility that interaction with FBPs is quantitatively affected.

Chemical shift index analysis of assigned residues of free human^38^ and *Dictyostelium*^9^ Skp1s indicated that the overall secondary structure elements of the dimers were similar to one another and to human Skp1 in complexes with FBPs, except for the C-terminal F-box subsite-2 region (residues 126-162) which was predominantly disordered in free Skp1’s. The current study of residues 1-125 extends to show that free Skp1 (dimer) assumes essentially the same structure as for human Skp1 bound to the FBP βTRCP, with an RMSD for the corresponding Cα atoms of 1.1 Å (Figure S6). Thus interactions with proteins including Cul1^8^ and Sgt1,^45^ whose crystallographically defined binding interfaces lie within this region but N-terminal to the dimer interface (ca. residues 1-90), are likely to be unaffected by the dimer status of Skp1.

Owing to the semi-parallel arrangement of the monomers, the two C-termini of the Skp1ΔΔ dimer are close enough to one another that the missing C-terminal regions have the potential to influence one another in the native protein. The availability of a stable monomeric form of Skp1 will now allow a direct NMR analysis of full length Skp1 and the consequences of its glycosylation, which has been postulated to influence the organization of F-box binding subsite-2^4,46^ – the C-terminal region that was truncated to enable the dimer structure reported here.

## Supporting information

Full supplement: tables S1-S2, Figs. S1-S8, additional methods

## ASSOCIATED CONTENT

### Supporting Information

The Supplement PDF contains Tables S1-S2, Figures S1-S8, and supplemental methods.

## Author Contributions

HWK and AE conducted the experiments. HWK, AE and KJG performed calculations. AE, HWK, EMS, JHP and CMW advised on design and interpretation. HWK, AE and CMW wrote the initial manuscript draft, which was edited by all authors and finalized by CMW. All authors approved the final version. ^‡^These authors contributed equally.

## Notes

The authors declare no competing financial interests.

## ACCESSION CODES

*Dictyostelium discoideum* Skp1A sequence: UniProtKB/Swiss-Prot: P52285.1

Skp1ΔΔ dimer NMR structure statistics: PDB ID: 6V88, BMRB ID:30696

## ACKNOWLEDGMENT

The authors thank John Glushka for NMR technical support.

## ABBREVIATIONS

AUC: analytical ultracentrifugation
FBP: F-box protein
HNOE: heteronuclear NOE
HSQC: heteronuclear single quantum coherence spectroscopy
NOESY: nuclear Overhauser enhancement spectroscopy
SCF: Skp1/cullin-1/F-box protein family of E3 ubiquitin ligases
Skp1Δ: Skp1Δinternal loop
Skp1ΔΔ: Skp1Δinternal loop/ΔC-terminus
TEV: tobacco etch virus

