## Supplementary material for "Skp1 Dimerization Conceals its F-box Protein Binding Site": Full supplement: tables S1-S2, Figs. S1-S8, additional methods

**SUPPLMENTARY INFORMATION**

### Table of Contents

Table S1. List of NMR experiments
Table S2. List of atom (or pseudoatom) pairs subject to 5 Å intermolecular distance constraints

Figure S1. Sequence of synthetic codon-optimized *Dictyostelium* $His_6$Skp1A
Figure S2. Sedimentation velocity analysis of Skp1ΔΔ
Figure S3. Skp1ΔΔ $^{15}$N-$^{1}$H HSQC resonance assignments
Figure S4. Intermolecular $^{1}$H-$^{1}$H NOE contacts between Skp1 dimer subunits
Figure S5. Dimer interface is composed of a 4-helix bundle
Figure S6. Overlay of a monomer from the Skp1ΔΔ dimer with human Skp1 from a crystal structure with an FBP
Figure S7. Heatmap of total free energy changes in the Skp1 monomer
Figure S8. Alignment of dimer interface sequences

Supplemental Methods

**SUPPLEMENTARY TABLES AND FIGURES**

**Table S1.** List of NMR experiments.

| Sample | Experiment* | Magnetic field (MHz) |
|---|---|---|
| Skp1ΔΔ | 2D [$^{15}$N, $^{1}$H]-HSQC | 800 |
| | 2D [$^{13}$C, $^{1}$H] CT-HSQC (aliphatic) | 800 |
| | 2D [$^{13}$C, $^{1}$H] CT-HSQC (aromatic) | 800 |
| | 3D $^{13}$C/$^{15}$N-edited [$^{1}$H,$^{1}$H] NOESY | 800 |
| | 3D TROSY-HNCACB | 800 |
| | 3D (H)CCH-COSY aromatic | 800 |
| | 3D (H)CCH-COSY aliphatic | 800 |
| | 3D (H)CCH-TOCSY aliphatic | 800 |
| | 3D $^{13}$C/$^{15}$N-filtered $^{13}$C/$^{15}$N-edited [$^{1}$H,$^{1}$H] NOESY | 600 |
| | 3D CBCA(CO)NH | 600 |
| | 3D HBHA(CO)NH | 600 |
| | 3D HNCO | 600 |
| | 3D (HCA)CONH | 600 |
| | 2D long-range [$^{15}$N, $^{1}$H]-HSQC | 600 |
| 100 uM Skp1ΔF97E | 2D [$^{15}$N, $^{1}$H]-HSQC | 900 |
| | 1D $^{15}$N $T_1$ | 600 |
| | 1D $^{15}$N $T_2$ | 600 |
| 500 uM Skp1ΔF97E | 2D [$^{15}$N, $^{1}$H]-HSQC | 900 |
| | 1D $^{15}$N $T_1$ | 600 |
| | 1D $^{15}$N $T_2$ | 600 |

* wiki.nesg.org

**Table S2.** List of atom (or pseudoatom) pairs subject to 5 Å intermolecular distance constraints. The constraints were used to ensure proper dimer formation during initial stages of structure calculation using CYANA.

| Atom 1 | Atom 2 |
|---|---|
| Phe97 HD | Ile100 HG2 |
| Phe97 HD | Ile100 HD1 |
| Phe97 HE | Ile100 HG2 |
| Phe97 HE | Ile100 HD1 |
| Phe97 HD | Val119 HG1 |
| Phe97 HD | Val119 HG2 |
| Phe97 HE | Val119 HG1 |
| Phe97 HE | Val119 HG2 |
| Phe97 HZ | Val119 HG1 |
| Phe97 HZ | Val119 HG2 |
| Phe97 HD | Met122 HE |
| Phe97 HE | Met122 HE |
| Phe97 HD | Ile123 HG2 |
| Phe97 HD | Ile123 HD1 |
| Phe97 HE | Ile123 HG2 |
| Phe97 HE | Ile123 HD1 |
| Leu101 HD1 | Lys117 HE |
| Leu101 HD2 | Lys117 HE |
| Leu101 HD1 | Ala120 HN |
| Leu101 HD2 | Ala120 HN |
| Leu101 HD1 | Ala120 HB |
| Leu101 HD2 | Ala120 HB |

**Figure S1.** Synthetic *Dictyostelium* Skp1A cDNAs. **(A)** The native (Dd) and synthetic (Syn) coding sequences of Skp1A are shown together with their identical translated amino acid sequences. Amino acid (bold) and nucleotide numbering starts at the original start Met. The start codons are highlighted in green, the new N-terminus (Ser) generated by TEV protease cleavage and Phe97 are highlighted in yellow, nucleotide replacements in the synthetic DNA are in red, and nucleotides 196-232 replaced in Skp1Δloop (panels B, C) are underlined. **(B)** The native (Dd) and synthetic (Syn) cDNA sequences for His$_6$Skp1Δloop, in which amino acids SPQGDDKKDEKR are replaced by GGSG (in blue). Original numbering for downstream positions is retained. Oligonucleotide sequences utilized for site-directed mutagenesis to generate F97E are indicated below the sequences, including the lower strand (AS) where needed to locate antisense primers. **(C)** Deletions were generated in the native sequence of Skp1A in pET19b-Skp1 by a variation of site-directed mutagenesis using the indicated oligonucleotides, with matching sequences highlighted in yellow or gray. Truncated C-terminal sequences are in light gray.

**(A)** His$_6$Skp1A

```
                                                       M  G  H   -20
Syn                                                  CCATGGgccac -60 NcoI

    H  H  H  H  H  S  S  G  V  D  L  G  T  E  N  L  Y  F  Q  S      0
Syn catcatcatcatcattcttctggtgtagatctgggtaccgagaacctgtacttccaatcc    0

    M  S  L  V  K  L  E  S  S  D  E  K  V  F  E  I  E  K  E  I     20
Dd  ATGTCTTTAGTTAAATTAGAATCTTCAGATGAAAAAGTCTTTGAAATTGAAAAAGAAATC   60
Syn ATGAGCCTGGTGAAACTGGAAAGCAGCGACGAGAAGGTGTTCGAGATTGAGAAAGAGATT

    A  C  M  S  V  T  I  K  N  M  I  E  D  I  G  E  S  D  S  P     40
Dd  GCTTGTATGTCAGTTACAATCAAGAATATGATTGAAGATATTGGTGAATCAGATAGTCCA  120
Syn GCGTGCATGAGCGTGACCATTAAAAACATGATCGAGGACATTGGTGAAAGCGATAGCCCG

    I  P  L  P  N  V  T  S  T  I  L  E  K  V  L  D  Y  C  R  H     60
Dd  ATTCCATTACCAAATGTTACTAGCACTATTTTAGAGAAAGTTCTTGACTATTGCAGACAT  180
Syn ATTCCGCTGCCGAACGTGACCAGCACCATCCTGGAGAAGGTTCTGGACTACTGCCGTCAC

    H  H  Q  H  P  S  P  Q  G  D  D  K  K  D  E  K  R  L  D  D     80
Dd  CACCATCAACATCCATCACCACAAGGTGACGATAAAAAGGATGAAAAGAGATTAGATGAT  240
Syn CATCACCAGCACCCGAGCCCGCAAGGTGACGATAAGAAAGATGAAAAGCGTCTGGACGAT

    I  P  P  Y  D  R  D  F  C  K  V  D  Q  P  T  L  F  E  L  I    100
Dd  ATCCCACCATATGATAGAGATTTCTGTAAAGTCGATCAACCAACCTTATTCGAATTAATC  300
Syn ATTCCGCCGTACGACCGTGATTTCTGCAAAGTGGACCAGCCGACCCTGTTTGAGCTGATT

    L  A  A  N  Y  L  D  I  K  P  L  L  D  V  T  C  K  T  V  A    120
Dd  TTGGCAGCCAATTATTTGGATATCAAACCATTATTAGATGTTACCTGTAAAACTGTTGCC  360
Syn CTGGCGGCGAACTATCTGGACATCAAGCCGCTGCTGGATGTGACCTGCAAAACCGTTGCG

    N  M  I  R  G  K  T  P  E  E  I  R  K  I  F  N  I  K  N  D    140
Dd  AATATGATCAGAGGTAAAACCCCAGAAGAAATCAGAAAAATCTTCAACATCAAGAACGAC  420
Syn AACATGATCCGTGGTAAAACCCCGGAGGAAATCCGTAAGATTTTCAACATCAAGAACGAT

    F  T  P  E  E  E  E  Q  I  R  K  E  N  E  W  C  E  D  K  G    160
Dd  TTTACTCCAGAAGAAGAAGAACAAATCAGAAAAGAAAATGAATGGTGTGAAGATAAAGGT  480
Syn TTCACCCCGGAAGAAGAGGAACAAATCCGTAAGGAGAACGAATGGTGCGAGGACAAGGGT

    G  N  *                                                        162
Dd  GGAAACTAA                                                      489
Syn GGTAACTAAGGATCC                                                BamHI
```

**(B)** His$_6$Skp1AΔloop

```
                                                       M  G  H    -20
Syn                                                 CCATGGgccac   -60 NcoI

    H  H  H  H  H  S  S  G  V  D  L  G  T  E  N  L  Y  F  Q  S       0
Syn catcatcatcatcattcttctggtgtagatctgggtaccgagaacctgtacttccaatcc     0

    M  S  L  V  K  L  E  S  S  D  E  K  V  F  E  I  E  K  E  I      20
Dd  ATGTCTTTAGTTAAATTAGAATCTTCAGATGAAAAAGTCTTTGAAATTGAAAAAGAAATC    60
Syn ATGAGCCTGGTGAAACTGGAAAGCAGCGACGAGAAGGTGTTCGAGATTGAGAAAGAGATT

    A  C  M  S  V  T  I  K  N  M  I  E  D  I  G  E  S  D  S  P      40
Dd  GCTTGTATGTCAGTTACAATCAAGAATATGATTGAAGATATTGGTGAATCAGATAGTCCA   120
Syn GCGTGCATGAGCGTGACCATTAAAAACATGATCGAGGACATTGGTGAAAGCGATAGCCCG

    I  P  L  P  N  V  T  S  T  I  L  E  K  V  L  D  Y  C  R  H      60
Dd  ATTCCATTACCAAATGTTACTAGCACTATTTTAGAGAAAGTTCTTGACTATTGCAGACAT   180
Syn ATTCCGCTGCCGAACGTGACCAGCACCATCCTGGAGAAGGTTCTGGACTACTGCCGTCAC

    H  H  Q  H  P  G  G  S  G                          L  D  D      80
Dd  CACCATCAACATCCAGGTGGTTCCGGA------------------------TTAGATGAT   240
Syn CATCACCAGCACCCGGGTGGTTCCGGA------------------------CTGGACGAT

    I  P  P  Y  D  R  D  F  C  K  V  D  Q  P  T  L  F  E  L  I     100
Dd  ATCCCACCATATGATAGAGATTTCTGTAAAGTCGATCAACCAACCTTATTCGAATTAATC   300
Syn ATTCCGCCGTACGACCGTGATTTCTGCAAAGTGGACCAGCCGACCCTGTTTGAGCTGATT    F97E-S
 AS TAAGGCGGCATGCTGGCACTAAAGACGTTTCACCTGGTCGGCTGGGACAAACTCGACTAA    F97E-AS

    L  A  A  N  Y  L  D  I  K  P  L  L  D  V  T  C  K  T  V  A     120
Dd  TTGGCAGCCAATTATTTGGATATCAAACCATTATTAGATGTTACCTGTAAAACTGTTGCC   360
Syn CTGGCGGCGAACTATCTGGACATCAAGCCGCTGCTGGATGTGACCTGCAAAACCGTTGCG    F97E-S
 AS GACCGCCGCTTGATAGACCTGTAGTTCGGCGACGACCTACACTGGACGTTTTGGCAACGC

    N  M  I  R  G  K  T  P  E  E  I  R  K  I  F  N  I  K  N  D     140
Dd  AATATGATCAGAGGTAAAACCCCAGAAGAAATCAGAAAAATCTTCAACATCAAGAACGAC   420
Syn AACATGATCCGTGGTAAAACCCCGGAGGAAATCCGTAAGATTTTCAACATCAAGAACGAT

    F  T  P  E  E  E  E  Q  I  R  K  E  N  E  W  C  E  D  K  G     160
Dd  TTTACTCCAGAAGAAGAAGAACAAATCAGAAAAGAAAATGAATGGTGTGAAGATAAAGGT   480
Syn TTCACCCCGGAAGAAGAGGAACAAATCCGTAAGGAGAACGAATGGTGCGAGGACAAGGGT

    G  N  *                                                        162
Dd  GGAAACTAA                                                      489
Syn GGTAACTAAGGATCC                                                BamHI
```

**F97E mutation:**

Dd-Skp1A-optim-F97E-S:  5’-GACCCTCGAGGAGCTGATTCTGGCGGCG

Dd-Skp1A-optim-F97E-AS: 5’-CAGCTCCTCGAGGGTCGGCTGGTCCAC

**(C)** Skp1AΔloopΔCterm (Skp1ΔΔ)

```
M   S   L   V   K   L   E   S   S   D   E   K   V   F   E   I   E   K   E   I
ATGTCTTTAGTTAAATTAGAATCTTCAGATGAAAAAGTCTTTGAAATTGAAAAAGAAATC

A   C   M   S   V   T   I   K   N   M   I   E   D   I   G   E   S   D   S   P
GCTTGTATGTCAGTTACAATCAAGAATATGATTGAAGATATTGGTGAATCAGATAGTCCA

I   P   L   P   N   V   T   S   T   I   L   E   K   V   L   D   Y   C   R   H
ATTCCATTACCAAATGTTACTAGCACTATTTTAGAGAAAGTTCTTGACTATTGCAGACAT
TAAGGTAATGGTTTACAATGATCGTGATAAAATCTCTTTCAAGAACTGATAACGTCTGTA  Δloop-AS

H   H   Q   H   P   G   G   S   G                                       L   D   D
CACCATCAACATCCAGGTGGTTCCGGA------------------------TTAGATGAT  Δloop-S
GTGGTAGTTGTAGGTAGTGGTGTTCCA------------------------AATCTACTA  Δloop-AS

I   P   P   Y   D   R   D   F   C   K   V   D   Q   P   T   L   F   E   L   I
ATCCCACCATATGATAGAGATTTCTGTAAAGTCGATCAACCAACCTTATTCGAATTAATC  Δloop-S
TAGGGTGGTATACTATCTCTAAAGACATTTCAGCTAGTTGGTTGGAATAAGCTTAATTAG

L   A   A   N   Y   L   D   I   K   P   L   L   D   V   T   C   K   T   V   A
TTGGCAGCCAATTATTTGGATATCAAACCATTATTAGATGTTACCTGTAAAACTGTTGCC
AACCGTCGGTTAATAAACCTATAGTTTGGTAATAATCTACAATGGACATTTTGACAACGG  ΔCterm-AS

N   M   I   R   G   K   T   P   E   E   I   R   K   I   F   N   I   K   N   D
AATATGATCAGAGGTAAAACCCCAGAAGAAATCAGAAAAATCTTCAACATCAAGAACGAC  ΔCterm-S
TTATACTAGTCTCCA                                               ΔCterm-AS

 F   T   P   E   E   E   E   Q   I   R   K   E   N   E   W   C   E   D   K   G
TTTACTCCAGAAGAAGAAGAACAAATCAGAAAAGAAAATGAATGGTGTGAAGATAAAGGT

G   N   *
GGAAACTAAGGATCCGGCTGCTAACAAAGCCCGAAAGGAAGCTGAGTTGGCTGCTGCCAC  ΔCterm-S
      ATTCCTAGG
```

**Deletions:**

```
ΔCterm-S: 5'-ATCAGAGGTTAAGGATCCGGCTGCTAACAAAGC
ΔCterm-AS:GGATCCTTAACCTCTGATCATATTGGCAACAGTTTTAC

Δloop-S: 5'-CAACATCCAGGTGGTTCCGGATTAGATGATATCCCACCATATGATAGAGATTTC
Δloop-AS: 5'-ATCATCTAATCCGGAACCACCTGGATGTTGATGGTGATGTCTGC
```

**Figure S2.** Sedimentation velocity analysis of Skp1ΔΔ. Analytical ultracentrifugation of 100 µM Skp1ΔΔ, as described in Figure 1. DdSkp1ΔΔ sediments at 1.92 S, which matches the predicted 1.92 value for the dimer using HYDROPRO. In contrast, the sedimentation coefficient for a monomeric Skp1ΔΔ was predicted be 1.28 S. The breadth of the peak width suggests that a small amount of monomeric Skp1ΔΔ may exist in rapid equilibrium with the dimer.

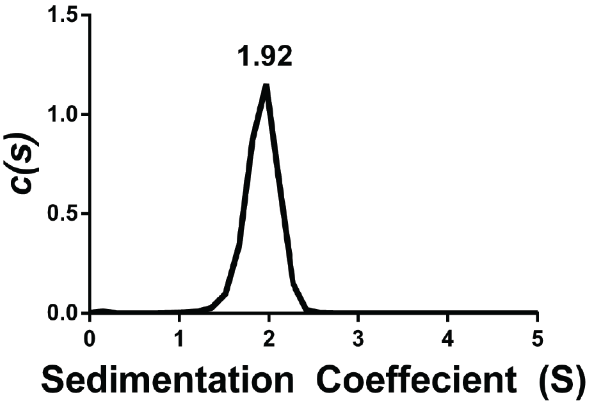

**Figure S3.** 2D [$^{15}$N, $^{1}$H] HSQC spectrum of a 1:1 mixed sample of U-$^{15}$N,$^{13}$C labeled and natural abundance Skp1ΔΔ acquired at 800 MHz field strength. Peak assignments are indicated in green with residue numbers corresponding to the wild-type sequence. Assignments associated with the non-native tetrapeptide insertion (GGSG) that replaced the internal loop (residues 66-77) are denoted with asterisks.

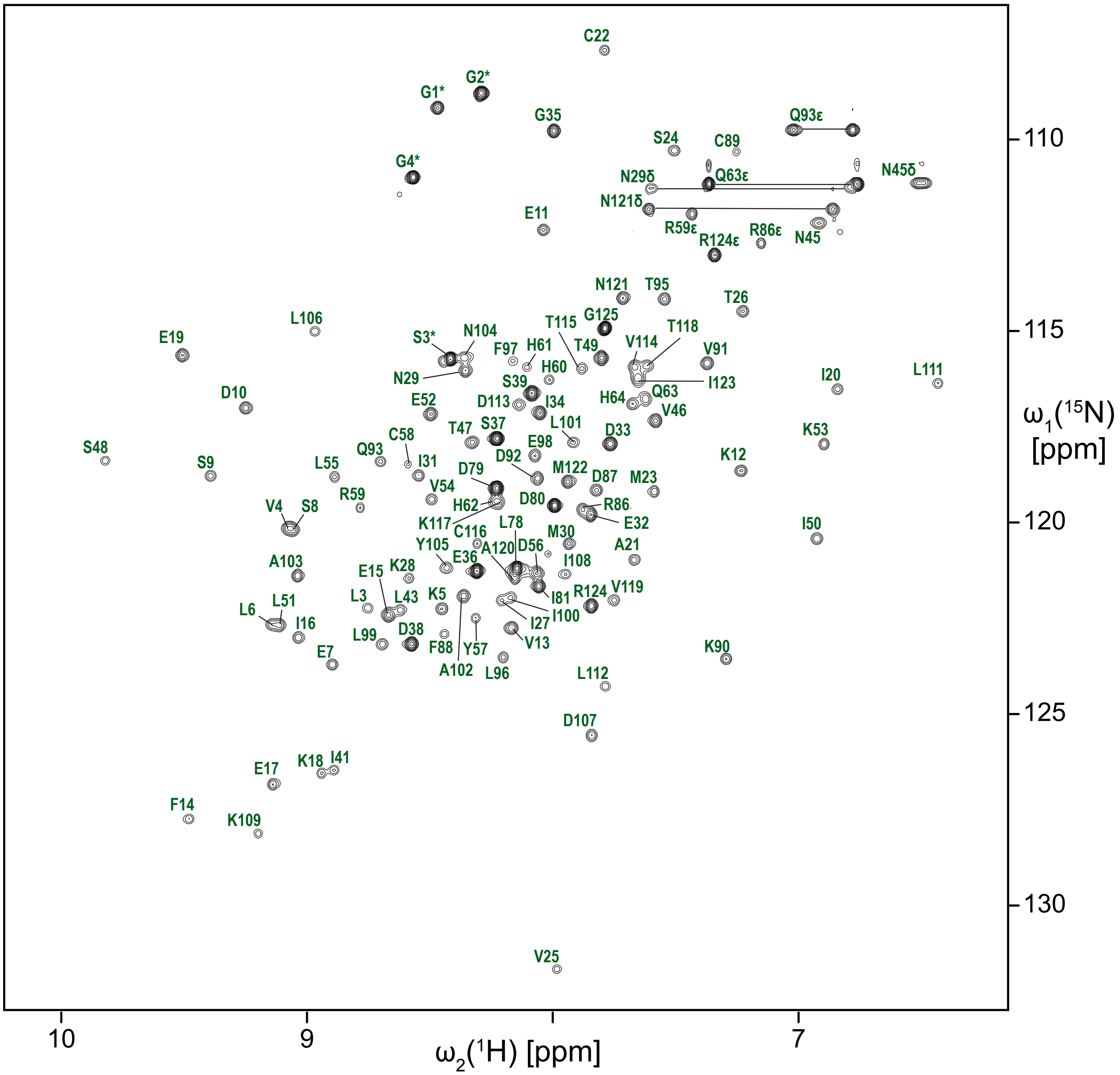

**Figure S4.** Intermolecular $^{1}H$-$^{1}H$ NOE contacts between Skp1ΔΔ dimer subunits. Representative strips from 3D $^{13}C$/$^{15}N$-edited [$^{1}H$,$^{1}H$] NOESY and 3D $^{13}C$/$^{15}N$-filtered, $^{13}C$-edited [$^{1}H$,$^{1}H$] NOESY spectra showing contacts for HD1 of L101 recorded on a mixture of natural abundance and $^{13}C$,$^{15}N$-enriched Skp1ΔΔ. Spectra were recorded at 800 and 600 MHz magnetic field strengths, with mixing times of 70 and 120 ms, respectively. Intermolecular and intramolecular NOE correlations are labeled in red and black, respectively.

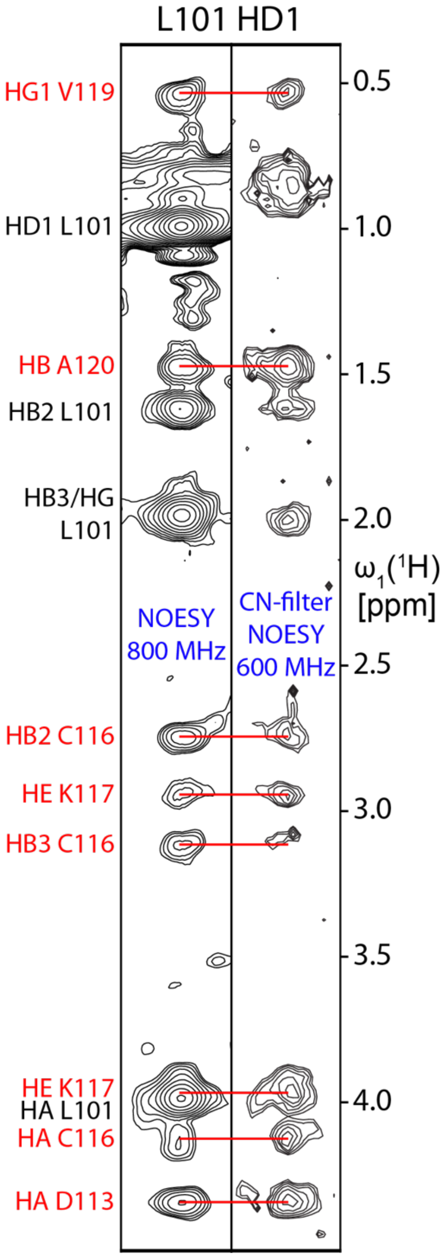

**Figure S5.** The dimer interface is composed of a 4-helix bundle, or a pair of pairs that are related by two-fold rotational symmetry. Green and magenta represent the helical pairs contributed by each Skp1ΔΔ subunit.

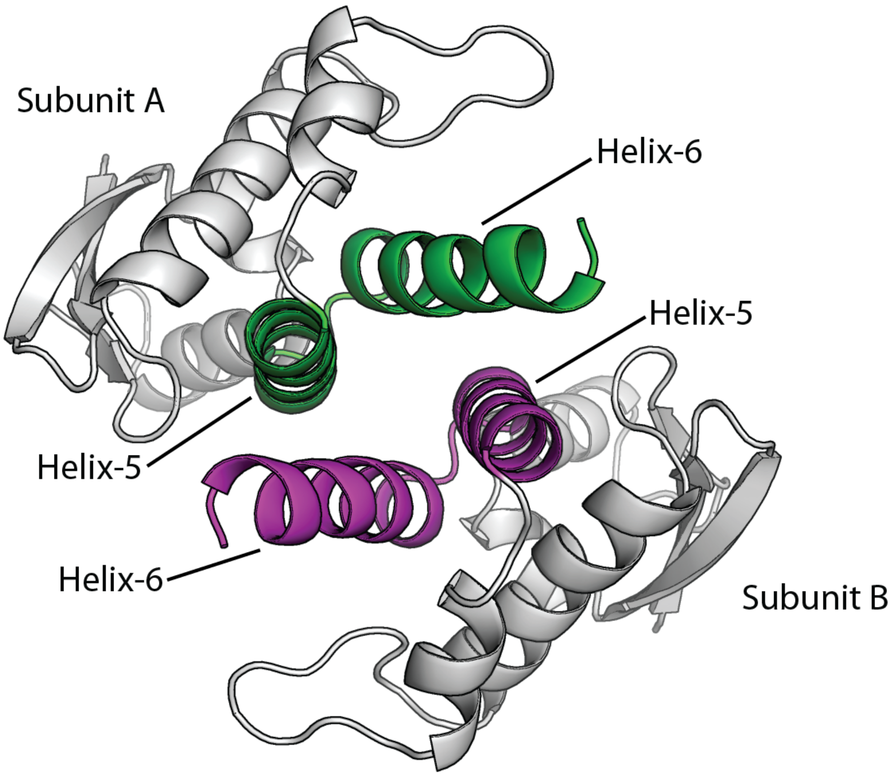

**Figure S6.** Overlay of Cα traces of a single subunit from the *Dictyostelium* Skp1ΔΔ dimer (green) and human Skp1 (red). HsSkp1 was excerpted from a crystal structure of its complex with βTRCP, a human FBP (PDB ID 6M90). The N- and C-termini are indicated. The corresponding 97 Cα atoms of DdSkp1 (amino acids 2-33, 40-62, 83-123) and HsSkp1 (amino acids 2-33, 38-60, 88-128) align with an RMSD of 1.1 Å. An internal loop in HsSkp1, which corresponds to the segment substituted with a GGSG tetrapeptide in DdSkp1ΔΔ, is absent from its Cα trace at the black triangle.

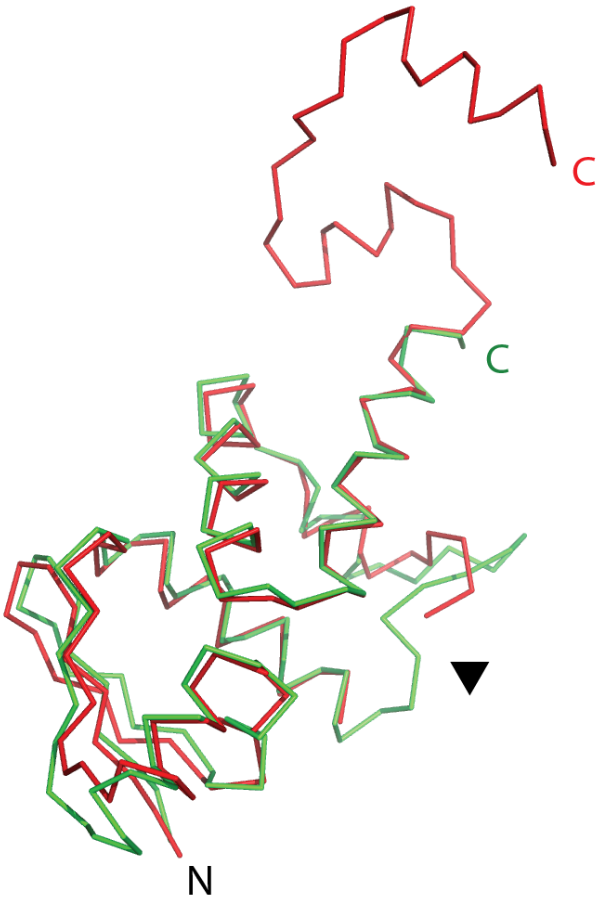

**Figure S7.** Heatmap of total *in silico* free energy changes in the Skp1 monomer for each amino acid substitution at the dimer interface. The effects of amino acid replacements are plotted on the y-axis as a for each interface position (x-axis). Colors represent calculated changes in the total free energy of the monomer (total ΔΔG score). Values greater than one (warmer colors) indicate destabilizing mutations, and values less than one (colder colors) imply stabilizing mutations.[31] Compare with effects on the dimer state (Fig. 3C).

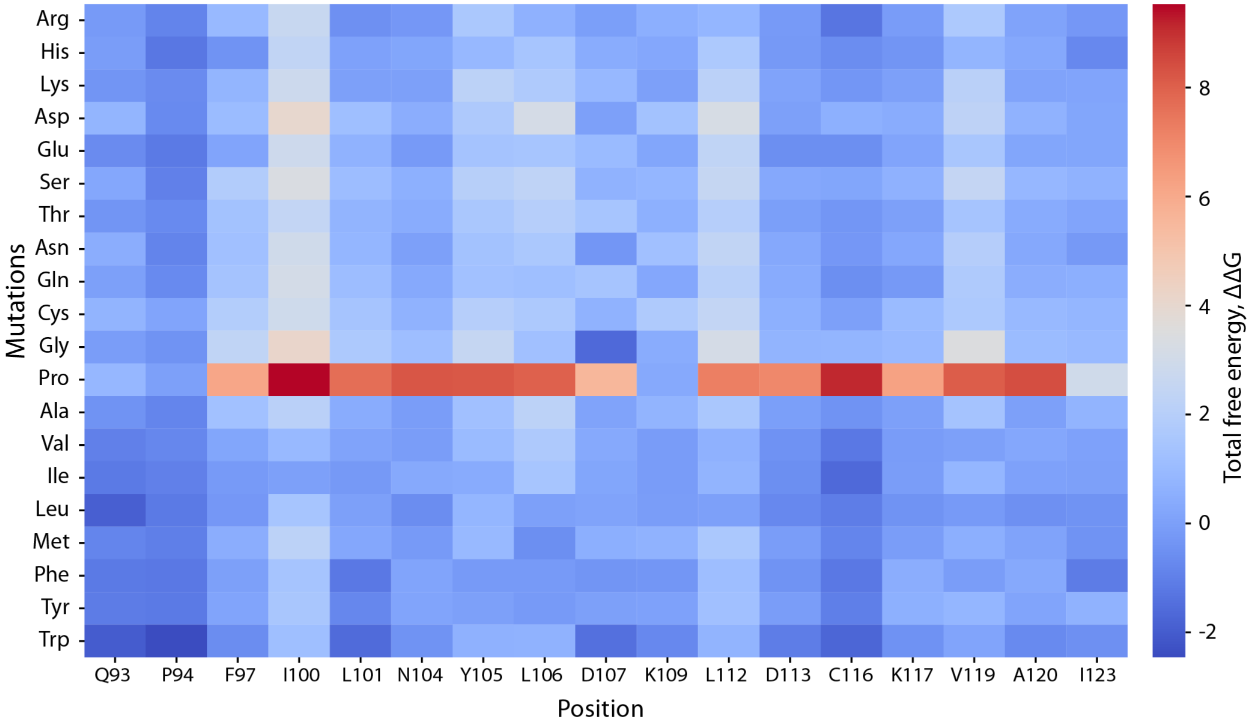

**Figure S8.** Alignment of Skp1 sequences from across eukaryotic phylogeny. The D letters in the top row indicate positions of intermolecular contacts observed for *Dictyostelium* Skp1ΔΔ. Lower case h letters depict the positions of helices-5, -6, -7, and -8 (from left to right) in crystal structures with FBPs. To facilitate visualization of relatedness, acidic residues are in blue, basic in dark red, Gly and Pro in red, and hydrophobic in green, as previously described.[47] Positions possessing a consensus chemical characteristic are highlighted, in yellow (hydrophobic), gray (acidic), dark grey (basic), or teal (small). Positions of near perfect conservation are bolded. Asterisk denotes the C-terminus.

```
                D   D  D        D  DD DD DD
              hhhhhhhhhhhhh   hhhhhhhhhhhhh     hhhhhhh        hhhhhhhhhhhh
DdSkp1 90-VDQPTLFELILAANYLDIKPLLDVTCKTVANMIRGKTPEEIRKIFNIKNDFTPEEEEQIRKENEWCEDKGGN*
Tg        SDQKILFALILAANYLNIKPLLDLSVAKVATMIKAKTPEEIRRIFNIVNDFTPEEEAQVREENKWCEDA*
Tp        VEQVLLFELILAANYMDIKPLLDLTCATVASMIKGKTPEEIRKTFNIANDFSPEEEAQVREENKWCEEP*
Pu-1      IEQDILFELILAANYMDIKSLLDLACAKVASMIKGKTPQEIRETFNIVNDFTPEEESQIREENKWCEEA*
Tt        LEQ--LFEIILAANYLDIKSLLDLACAKVATLIKNKTPDEIRKTFNIPNDFTPEEEAQIREENKWAEEATS*
Cr        VDQATLFDLILAANYLNIKGLLDLTCQTVAQMIKGKTPEEIRKTFNIKNDFTPEEEEEVRRENQWAFD*
Sc        VDQEMLYEIILAANYLNIKPLLDAGCKVVAEMIRGRSPEEIRRTFNIVNDFTPEEEAAIRRENEWAEDR*
Ce-1      VDQGTLFELILAANYLDIKGLLDVTCKTVANMIKGKSPEEIRRTFNIKNDFTPEEEEQIRKENAWCED*
Hs        VDQGTLFELILAANYLDIKGLLDVTCKTVANMIKGKTPEEIRKTFNIKNDFTEEEEAQVRKENQWCEEK*
```

Dd, *Dictyostelium discoideum* (cellular slime mold), NCBI: AAA67888.1
Tg, *Toxoplasma gondii* (apicomplexa), ToxoDB: TGARI_207680
Tp, *Thalassiosira pseudonana* 38460 (diatom/stramenopile), NCBI, XP_002294707.1
Pu, *Pythium ultimum*, oomycete plant pathogen, PYU1_T013675, www.pythiumdb.org
Tt, *tetrahymena thermophila* (ciliated protozoan), TTHERM_00426320
Cr, *Chlamydomonas reinhardtii* (green algae/chlorophyta), XP_001690964.1
Sc, *Saccharomyces cerevisiae* (budding yeast), NCBI, NP_010615.3
Ce, *Caenorhabditis elegans* (nematoda) NP_492513.1
Hs, *Homo sapiens* (vertebrata), NCBI, NP_008861.2

### SUPPLEMENTAL METHODS

The scripts below record the methods employed to analyze the Skp1 dimer structure.

- ***FastRelax protocol***

  The FastRelax protocol used for structure preparation is shown below:

```
<ROSETTASCRIPTS>
    <SCOREFXNS>
            <ScoreFunction name="ref2015" weights="ref2015" symmetric="1">
                    <Reweight scoretype="atom_pair_constraint" weight="1.0"/>
            </ScoreFunction>
    </SCOREFXNS>
    <MOVERS>
            <SetupForSymmetry name="setupsymm" definition="Skp1.symm"/>
            <AddConstraintsToCurrentConformationMover name="pair_cst" use_distance_cst="1"/>
            <FastRelax name="fast_relax" scorefxn="ref2015" repeats="3">
                    <MoveMap name="moveMap" bb="1" chi="1" jump="1"/>
            </FastRelax>
    </MOVERS>
    <PROTOCOLS>
            <Add mover="setupsymm"/>
            <Add mover="pair_cst"/>
            <Add mover="fast_relax"/>
    </PROTOCOLS>
    <OUTPUT scorefxn="ref2015"/>
</ROSETTASCRIPTS>
```

  This protocol was run using as input files the protein structure (pdb file) and a symmetry definition file called Skp1.symm. The symmetry definition file was generated with a public script (available at $Rosetta/main/source/src/apps/public/symmetry/make_symmdef_file.pl), using the command line options -m NCS -a A -i B -p input.pdb > Skp1.symm.

- ***Flex ddG protocol***

  The flex ddG protocol was implemented as described by Barlow et al. (2018). The corresponding script can be found at https://github.com/Kortemme-Lab/flex_ddG_tutorial/blob/master/ddG-backrub.xml.
  The command line options used to run the protocol were:

  $Rosetta/main/source/bin/rosetta_scripts.linuxgccrelease
  - s refined_Skp1.pdb
  - parser:protocol flexddg_protocol.xml
  - parser:script_vars chainstomove=B mutate_resfile=mutation_file number_backrub_trials=35000 max_minimization_iter= 5000 abs_score_convergence_thresh=10 backrub_trajectory_stride=35000

- restore_talaris_behavior
- in:file:fullatom
- ignore_unrecognized_res
- ignore_zero_occupancy false
- ex1 ex2
- nstruct 35

- ***Cartesian_ddG protocol***

  Prior to the mutagenesis simulation, the refined Skp1 dimer was relaxed in Cartesian space using the command line options:

```
$Rosetta/main/source/bin/relax.linuxgccrelease
-database $Rosetta/main/database
-s refined_Skp1.pdb
-use_input_sc
-constrain_relax_to_start_coords
-ignore_unrecognized_res
-nstruct 15
-relax:coord_constrain_sidechains
-relax:cartesian
-score:weights ref2015_cart
-relax:min_type lbfgs_armijo_nonmonotone
-relax:script cart2.script
-symmetry_definition skp1.symm
```

  With the cart2.script file:

```
switch:cartesian
repeat 2
ramp_repack_min 0.02  0.01     1.0  50
ramp_repack_min 0.250 0.01     0.5  50
ramp_repack_min 0.550 0.01     0.0 100
ramp_repack_min 1     0.00001  0.0 200
accept_to_best
endrepeat
```

  The Cartesian_ddg protocol was used as described by Park et al. (2016). The corresponding script can be found at https://www.rosettacommons.org/docs/latest/cartesian-ddG.
